# Interplay between substrate recognition, 5’ end tRNA processing and methylation activity of human mitochondrial RNase P

**DOI:** 10.1101/452920

**Authors:** Agnes Karasik, Carol A. Fierke, Markos Koutmos

## Abstract

Human mitochondrial ribonuclease P (mtRNase P) is an essential three protein complex that catalyzes the 5’ end maturation of mitochondrial precursor tRNAs (pre-tRNAs). MRPP3 (Mitochondrial RNase P Protein 3), a protein-only RNase P (PRORP), is the nuclease component of the mtRNase P complex and requires a two-protein S-adenosyl methionine (SAM)-dependent methyltransferase MRPP1/2 sub-complex to function. Dysfunction of mtRNase P is linked to several human mitochondrial diseases, such as mitochondrial myopathies. Despite its central role in mitochondrial RNA processing, little is known about how the protein subunits of mtRNase P function synergistically. Here we use purified mtRNase P to demonstrate that mtRNase P recognizes, cleaves, and methylates some, but not all, mitochondrial pre-tRNAs *in vitro*. Additionally, mtRNase P does not process all mitochondrial pre-tRNAs uniformly, suggesting the possibility that some pre-tRNAs require additional factors to be cleaved *in vivo*. Consistent with this, we found that addition of the MRPP1 co-factor SAM enhances the ability of mtRNase P to bind and cleave some mitochondrial pre-tRNAs. Furthermore, the presence of MRPP3 can enhance the methylation activity of MRPP1/2. Taken together, our data demonstrate that the subunits of mtRNase P work together to efficiently recognize, process and methylate human mitochondrial pre-tRNAs.

## INTRODUCTION

Ribonuclease P (RNase P) catalyzes one of the first steps of tRNA maturation, the removal of the 5’ leader sequences from precursor tRNAs (pre-tRNAs). There are two classes of evolutionarily distinct RNase Ps: RNA-based (ribozyme) and protein-based enzymes (Gobert et al. 2010; Holzmann et al. 2008; Taschner et al. 2012). RNase P is essential wherever pre-tRNAs are transcribed and is present in the nucleus, mitochondria and chloroplasts of cells. The protein-only form of RNase P was first identified in human mitochondria and consists of three subunits: <u>M</u>itochondrial <u>R</u>Nase <u>P P</u>rotein 1 (MRPP1, TRMT10C), <u>M</u>itochondrial <u>R</u>Nase <u>P P</u>rotein 2 (MRPP2, 17β-hydroxysteroid dehydrogenase type 1, SDR5C1), and <u>M</u>itochondrial <u>R</u>Nase <u>P P</u>rotein 3 (MRPP3 or human PRORP, catalytic nuclease subunit) (Holzmann et al. 2008; Vilardo et al. 2012). Subsequently, <u>Pr</u>otein-<u>O</u>nly <u>R</u>Nase <u>Ps</u> (PRORPs) have been discovered in some land plants (e.g. *Arabidopsis thaliana* (Gobert et al. 2010; Gutmann, Gobert, and Giege 2012)) and other unicellular Eukaryotic organisms (e.g. *Trypanosoma bruceii* (Taschner et al. 2012)). It is noteworthy that in archaea and bacteria, 5’ tRNA enzymes containing a similar catalytic domain to PRORPs were also recently identified (HARPs, <u>H</u>omologs of <u>A</u>quifex <u>R</u>Nase <u>P</u>). These enzymes were shown to be the smallest RNase P with independent 5’ end pre-tRNA processing endonuclease activity (Nickel et al. 2017). In contrast to human mtRNase P, PRORPs are stand-alone enzymes that do not require accessory proteins. Most studies of protein-based RNase Ps have focused on PRORPs because these enzymes are homologous to MRPP3 and provide a tractable system for study (Pinker et al. 2017; Gobert et al. 2013; Gutmann, Gobert, and Giege 2012; Gobert et al. 2010; Karasik et al. 2016; Howard et al. 2013; Howard et al. 2012; Howard et al. 2016; Klemm et al. 2016). As a consequence, little is known about how the components of the metazoan mtRNase P complex function.

In mitochondria, RNAs are transcribed in polycistronic units. In each polycistronic unit, most protein-coding and rRNA transcripts are separated (or ‘punctuated’) by tRNAs. Therefore 5’ end and 3’ end tRNA processing is required for the processing of most key mitochondrial RNAs (Ojala, Montoya, and Attardi 1981). 5’ End cleavage by mtRNase P has been suggested to be the first step in the hierarchical tRNA processing in mitochondria (Sanchez et al. 2011; Reinhard, Sridhara, and Hallberg 2017; Rossmanith et al. 1995). More than 50% of mitochondrial disease-causing mutations in the mitochondrial genome are found in tRNA genes (Abbott, Francklyn, and Robey-Bond 2014). Since precursor tRNA processing is likely the first step in the mitochondrial tRNA maturation pathway, disabling this function leads to accumulation of mitochondrial precursor tRNAs along with unprocessed RNA transcripts (Sen et al. 2016; Rackham et al. 2016). Accumulation of unprocessed RNA in the mitochondria affects downstream processes such as mitochondrial ribosome assembly and translation with detrimental effects on mitochondrial function and viability. It is therefore predicted that mitochondrial RNA processing plays a role in mitochondrial diseases. Consistent with this idea, mtRNase P dysfunction is linked to several mitochondrial diseases including maternally inherited essential hypertension (Wang et al. 2011), mitochondrial myopathy (Bindoff et al. 1993; Rossmanith and Karwan 1998), MELAS (Li and Guan 2010), and HSD10-disease (Deutschmann et al. 2014; Chatfield et al. 2015).

All three subunits of mtRNase P are strictly required *in vivo*; knockout of any component in flies leads to larval lethality (Sen et al. 2016). In addition, knock-down of MRPP1 and MRPP3 in human cell lines was shown to affect 5’ end mitochondrial pre-tRNA processing and mature mitochondrial tRNA levels (Holzmann et al. 2008; Sanchez et al. 2011). This further emphasizes that in general, processing activity of MRPP3 is dependent on other components of mtRNase P. Similarly, knock-out of the MRPP3 gene in mice arrests embryogenesis at a very early stage (Rackham et al. 2016). Furthermore, mice containing a conditional knock-out of MRPP3 develop cardiomyopathy, show signs of mitochondrial dysfunction, and exhibit severe defects in 5’ end RNA processing (Rackham et al. 2016). MRPP3 has also been shown to further influence tRNA modifications; for example a missense mutation in MRPP3 was identified as an important factor responsible for differences in mitochondrial tRNA methylation patterns (Hodgkinson et al. 2014). Thus, correct mtRNase P function is likely to be essential for human health.

Little is known about the structure and function of the mtRNase P complex; there are no structures or structural models and limited mechanistic and biochemical information available, although structures of truncated MRPP3 have been solved (Karasik et al. 2016; Howard et al. 2012; Reinhard, Sridhara, and Hallberg 2015; Li et al. 2015). As such, the role of the interaction between MRPP3 and the MRPP1/2 sub-complex in enhancing substrate binding and catalysis is a key question in the function of human mtRNase P (Holzmann et al. 2008). The MRPP1/2 sub-complex has been predicted to methylate the N1 of adenine or guanine residues at the 9^th^ position of 19 mitochondrial tRNAs (Vilardo et al. 2012). MRPP1 is the S-adenosyl-methionine (SAM) dependent methyltransferase subunit in the MRPP1/2 sub-complex, and MRPP2 likely acts as a scaffold protein (Vilardo et al. 2012).

Here we investigate how MRPP1, MRPP2, and MRPP3 work together to recognize and cleave pre-tRNA substrates. Our work with recombinant mtRNase P reveals that the complex can specifically bind, process and methylate several human mitochondrial pre-tRNAs which are not recognized and processed by single-subunit plant PRORPs. Additionally, a subset of human mitochondrial pre-tRNA substrates are not processed efficiently by the recombinant mtRNase P complex suggesting that additional components or modifications may be required to facilitate cleavage of these substrates. We further demonstrate that the MRPP1 cofactor SAM enhances the ability of the mtRNase P complex to bind and cleave some substrates. Overall, our studies deepen insight into how the components of mtRNase P act synergistically to bind, process and methylate human mitochondrial pre-tRNAs.

## RESULTS

### mtRNase P binds mitochondrial precursor-tRNAs non-uniformly

Although the human mtRNase P complex was discovered almost a decade ago, it is largely unknown how it recognizes and binds its mitochondrial substrates (Holzmann et al. 2008). The pool of native mtRNase P substrates is diverse; in human mitochondria there are four subtypes of tRNAs (Type 0-III). The (mt)tRNAs belonging to types I, II, and III all differ from the canonical cloverleaf secondary structure of nuclear tRNAs (Helm et al. 2000; Suzuki, Nagao, and Suzuki 2011). We selected six mitochondrial pre-tRNA ((mt)pre-tRNAs) substrates to study representatives of all four classes including a substrate ((mt)pre-tRNA^Ser(AGY)^) that human mtRNase P is not expected process *in vivo* (Rossmanith 1997) (Figure 1). The exact lengths of 5’ and 3’ flanking ends of mtRNase P substrates have not yet been established; therefore, we used our knowledge of *A. thaliana* single subunit PRORP enzymes that are homologous to MRPP3 to design the 5’ and 3’ ends of our (mt)pre-tRNA substrates. PRORPs preferentially bind substrates with short (<5-8 nucleotide) 5’ leaders and do not bind the 3’ trailer of pre-tRNAs (Karasik et al. 2016; Howard et al. 2016; Brillante et al. 2016). Thus, the (mt)pre-tRNA substrates studied here possess 6-7 nucleotide long 5’ leader sequences and lack mature 3’ CCA ends (Figure 1).

**Figure 1.**
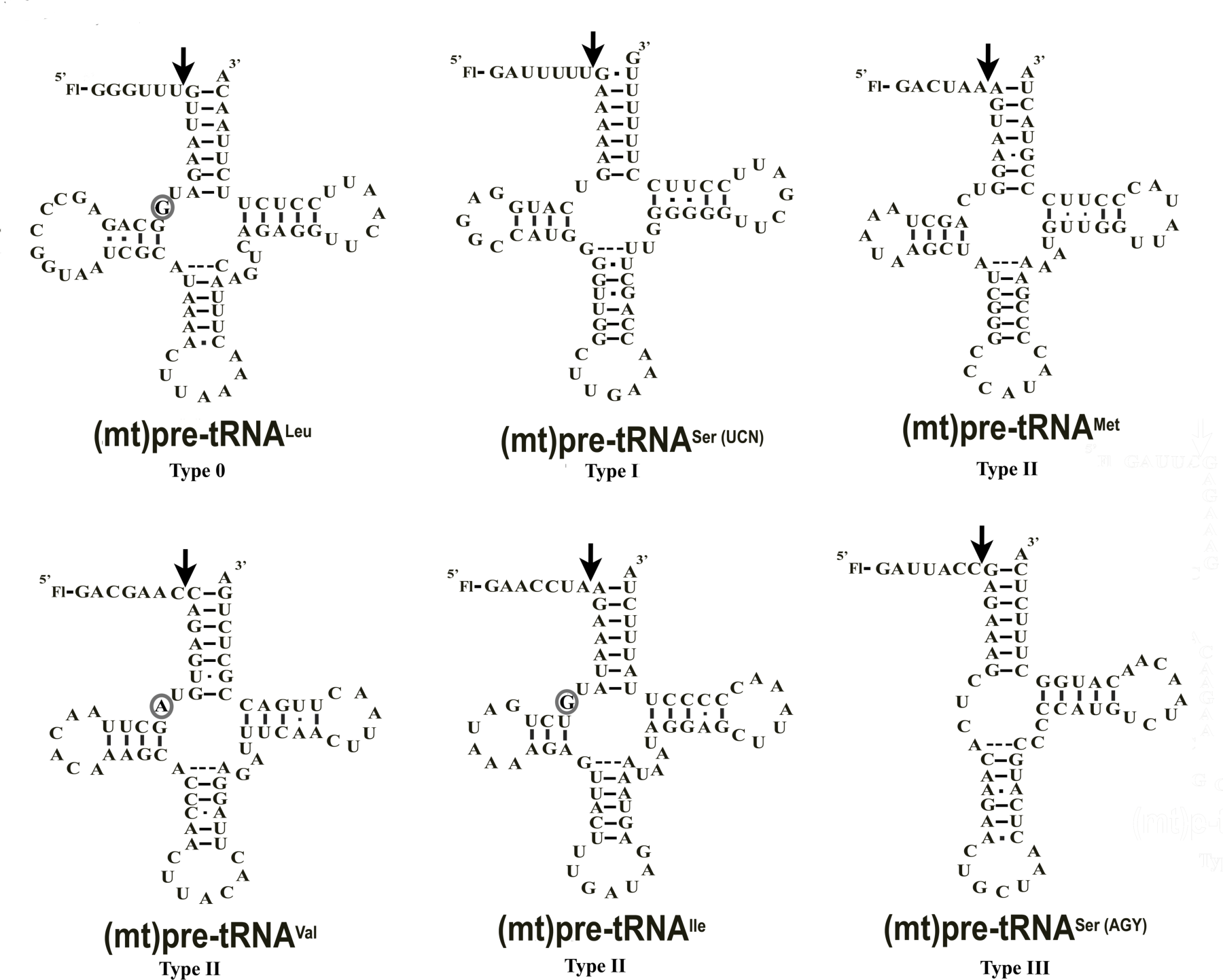
Proposed secondary structures of different types of human mitochondrial pre-tRNAs used in this study.

Black arrows indicate the 5’ cleavage site and ellipses mark the m1A9/m1G9 methylation sites for MRPP1/2.

To determine how each component of the mtRNase P complex interacts with (mt)pre-tRNAs we measured the binding constant (*K*_D_ or *K*_D,app_) for the mtRNaseP complex components (MRPP1/2 sub-complex and MRPP3) and six (mt)pre-tRNAs of interest using fluorescence anisotropy binding assay. We find that the isolated mtRNase P components generally do not associate tightly with the selected substrates (Table 1, Figure 2A and B). MRPP3 alone binds all of the (mt)pre-tRNAs substrates weakly with *K*_D_ values > 4 µM (Table 1 and Figure 2A). The MRPP1/2 sub-complex similarly binds most substrates weakly (*K*_D,app_ > 2 µM) with the exception of (mt)pre-tRNA^Ile^ and (mt)pre-tRNASer^UCN^, which it binds more tightly (*K*_D,app_ values of 104 ± 24 nM and 730 ± 196 nM, respectively) (Table 1, Figure 2B).

**Figure 2.**
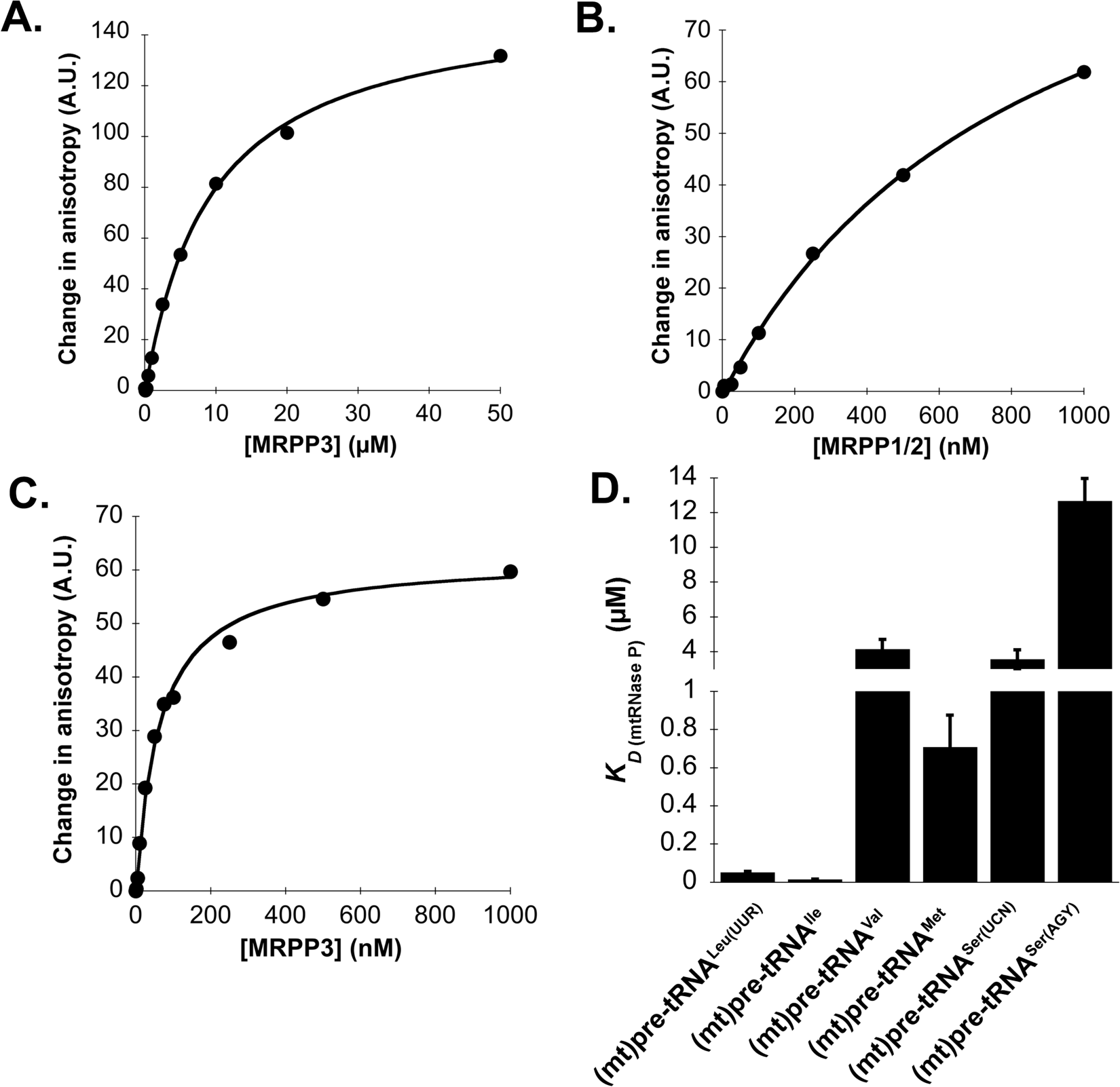
Presence of MRPP1/2 is required for efficient recognition of mitochondrial pre-tRNAs.

**A.** Representative figure of pre-tRNA binding to MRPP3 measured by changes in fluorescence polarization. 20 nM Fl-(mt)pre-tRNA^Leu(UUR)^ was titrated against increasing concentrations of MRPP3 (0-50 µM) using standard binding conditions. **B.** Representative figure of pre-tRNA binding to MRPP1/2 measured by changes in fluorescence polarization. 20 nM Fl-(mt)pre-tRNA^Leu(UUR)^ was titrated against increasing concentrations of MRPP1/2 (0-1 µM) using standard binding conditions. **C.** Representative figure of pre-tRNA binding to MRPP3 in the presence of 150 nM MRPP1/2 measured by changes in fluorescence polarization. 20 nM Fl-pre-(mt)tRNA^Leu(UUR)^ was titrated against increasing amount of MRPP3 (0-1 µM) using standard binding conditions. A.U.= arbitrary unit **D.** Average dissociation constants for the mtRNase P complex and investigated pre-tRNAs, measured as described in C.

**Table 1.**
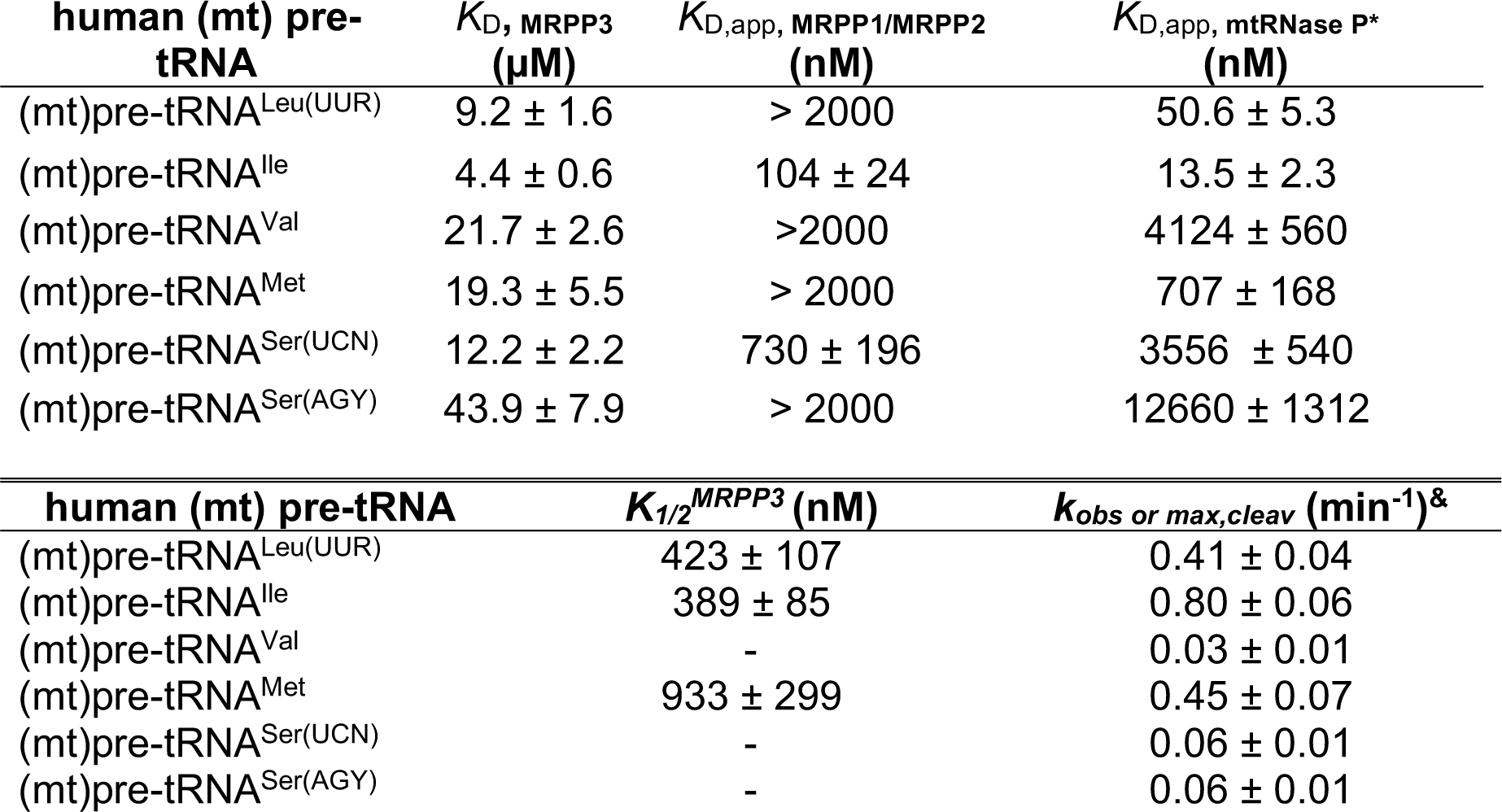
Human mtRNase P binds and processes a subset of mitochondrial pre-tRNA substrates efficiently.

| human (mt) pre-tRNA | $K_D$, MRPP3 (μM) | $K_{D,app}$, MRPP1/MRPP2 (nM) | $K_{D,app}$, mtRNase P* (nM) |
| --- | --- | --- | --- |
| (mt)pre-tRNA<sup>Leu(UUR)</sup> | 9.2 ± 1.6 | > 2000 | 50.6 ± 5.3 |
| (mt)pre-tRNA<sup>Ile</sup> | 4.4 ± 0.6 | 104 ± 24 | 13.5 ± 2.3 |
| (mt)pre-tRNA<sup>Val</sup> | 21.7 ± 2.6 | >2000 | 4124 ± 560 |
| (mt)pre-tRNA<sup>Met</sup> | 19.3 ± 5.5 | > 2000 | 707 ± 168 |
| (mt)pre-tRNA<sup>Ser(UCN)</sup> | 12.2 ± 2.2 | 730 ± 196 | 3556 ± 540 |
| (mt)pre-tRNA<sup>Ser(AGY)</sup> | 43.9 ± 7.9 | > 2000 | 12660 ± 1312 |

| human (mt) pre-tRNA | $K_{1/2}^{MRPP3}$ (nM) | $k_{obs}$ or $k_{max,cleav}$ (min<sup>-1</sup>)<sup>&</sup> |
| --- | --- | --- |
| (mt)pre-tRNA<sup>Leu(UUR)</sup> | 423 ± 107 | 0.41 ± 0.04 |
| (mt)pre-tRNA<sup>Ile</sup> | 389 ± 85 | 0.80 ± 0.06 |
| (mt)pre-tRNA<sup>Val</sup> | - | 0.03 ± 0.01 |
| (mt)pre-tRNA<sup>Met</sup> | 933 ± 299 | 0.45 ± 0.07 |
| (mt)pre-tRNA<sup>Ser(UCN)</sup> | - | 0.06 ± 0.01 |
| (mt)pre-tRNA<sup>Ser(AGY)</sup> | - | 0.06 ± 0.01 |

All fluorescence binding measurements were carried out in 30 mM MOPS pH=7.8, 150 mM NaCl, 1 mM DTT and 20 nM Fl pre-tRNAs. *K*_D_ values were calculated from a fit of a binding isotherm to the concentration dependence of the anisotropy data.

* Measurements for mtRNAse P were performed as following; first folded pre-tRNA was incubated with 150 nM MRPP1/2 for 5 minutes then varying amount of MRPP3 was added. Fluorescence anisotropy values were read after 5 minutes further incubation.

& These *k_max_* values were derived from *k_obs_* versus MRPP3 concentration plots for (mt)pre-tRNA^Ile^, (mt)pre-tRNA^Leu(UUR)^ and (mt)pre-tRNA^Met^. For all other (mt)pre-tRNAs data represent the *k_obs_* values using 500 nM MRPP1/2 and 1.5 µM MRPP3.

We next sought to establish how the fully-reconstituted mtRNase P complex (MRPP1/2:3) interacts with (mt)pre-tRNA substrates. For these assays, first we incubated pre-folded (mt)pre-tRNAs with 150 nM MRPP1/2. After MRPP1/2 and pre-tRNA binding reached equilibrium (5 minutes) we added varying concentrations of MRPP3 (0-10 ↃM, Figure 2C and D) and measured additional changes in fluorescence anisotropy. In general, we find that the presence of the MRPP1/2 sub-complex enhances (mt)pre-tRNA binding by 30 to 330-fold over binding to MRPP3 alone (Table 1). However, these enhancements are not uniform. The substrates fall into two categories: those that bind the complex somewhat tighter than MRPP3 alone (3-5 fold: (mt)pre-tRNA^Val^, (mt)pre-tRNASer^UCN^, (mt)pre-tRNA^Ser(AGY)^) and those that bind the complex significantly tighter (30-330 fold: (mt)pre-tRNA^Leu^, (mt)pre-tRNA^Ile^, (mt)pre-tRNA^Met^). As anticipated the substrate that binds the weakest is (mt)pre-tRNA^Ser(AGY)^. Notably, the mtRNase P complex exhibits the tightest *K*_D,app_ (13.5 ± 2.3 nM) for (mt)pre-tRNA^Ile^. These results indicate that while a subset of human (mt)pre-tRNAs are recognized with high affinity by mtRNase P (*K*_D,app_ values are in the nanomolar range), there are other (mt)pre-tRNAs that have weaker affinity (*K*_D,app_ values in the micromolar range) and may require additional factors or modifications to bind tightly to mtRNase P.

To compare substrate selectivity of human mtRNase P complex with single component PRORP1 and 2 enzymes we tested the ability of *A. thaliana* PRORP1 and PRORP2 to bind our six human (mt)pre-tRNA substrates (Table 1 and Supplemental Table 2). We find that only (mt)pre-tRNA^Ser(UCN)^ binds to PRORP1 and 2 with a comparable *K*_D_ value (60-120 nM) to that of their native *A. thaliana* substrates. All other (mt)pre-tRNAs were bound at least 7-fold less tightly (*K*_D_ = 880 nM to > 10 µM, Supplemental Table 2). Thus, mtRNase P uses strategies or interactions for recognizing its substrates that are different from the single-component PRORP1 and 2 enzymes.

### mtRNase P exhibits 5’ end processing of tightly bound (mt)pre-tRNAs

To further understand mtRNaseP substrate selectivity we measured single turnover cleavage rate constants for the six (mt)pre-tRNAs discussed above (Figure 1). Previously, we used fluorescence polarization-based multiple turnover assays for monitoring pre-tRNA 5’ end cleavage by RNA-based and single-component protein only RNase Ps (*A. thaliana* PRORPs)(Liu, Chen, and Fierke 2014; Howard et al. 2016). Here, we adapted this method to monitor the cleavage of 5’-fluorescein labeled (mt)pre-tRNAs by the fully-reconstituted human mtRNase P complex in single turnover cleavage assays. We measured the single turnover rate constants (*k*_obs_) and 
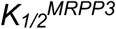
 values for mtRNaseP using limiting substrate (20 nM fluorescent pre-tRNA) and a range (100 nM-2 µM) of MRPP3 concentrations at 28° C while MRPP1/2 concentration was maintained constant at near saturating levels (500 nM) (Figure 3A and B, Supplemental Figure 3B).

**Figure 3.**
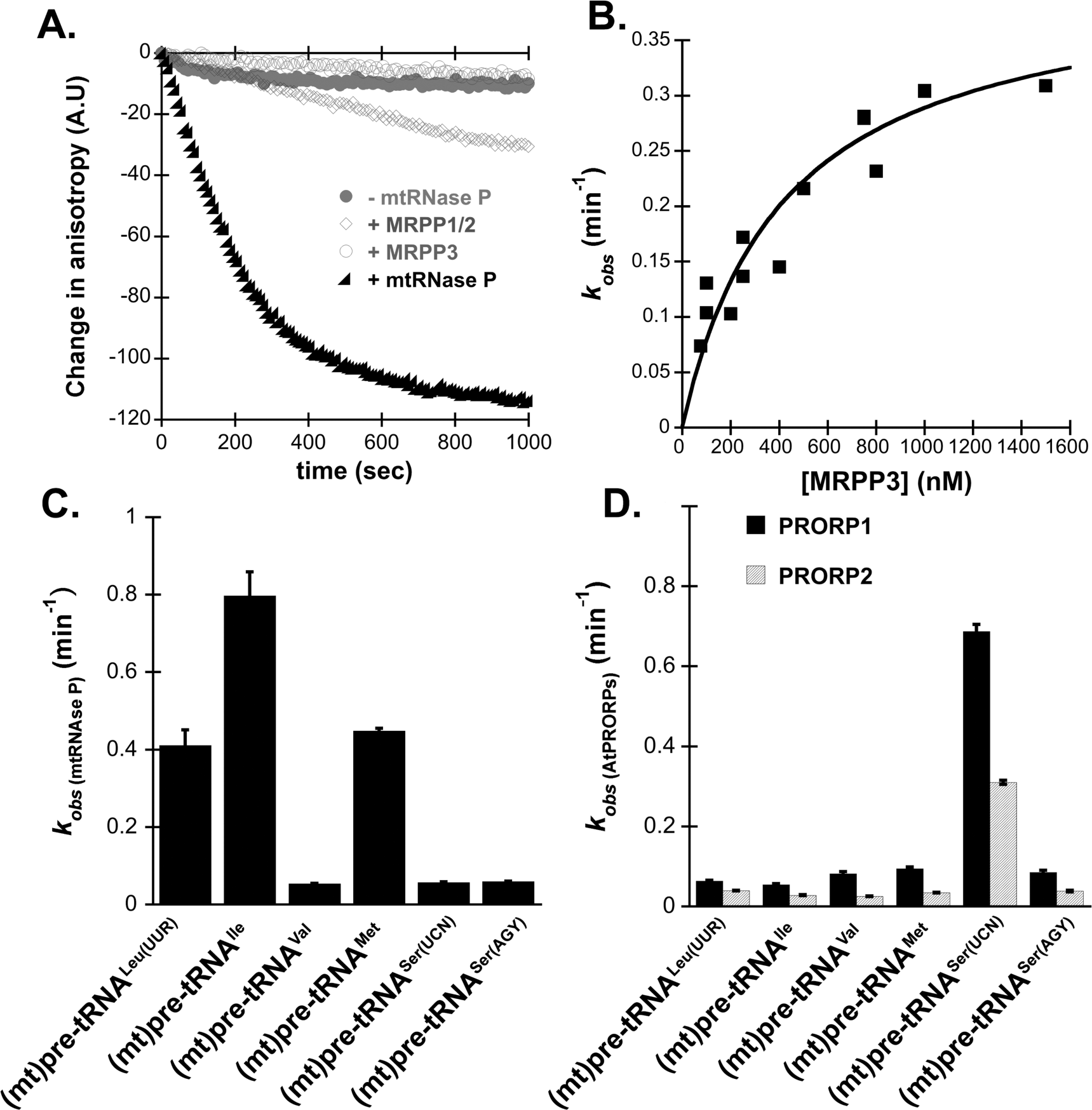
Some human mitochondrial pre-tRNAs are efficiently cleaved by the mtRNase P complex *in vitro.*

**A.** Representative figure of single turnover cleavage of 20 nM (mt)pre-tRNA^Leu(UUR)^ measured using changes in fluorescence polarization in the presence of the mtRNase P complex (closed triangles), MRPP3 only (open circles), MRPP1/2 only (open diamonds). Control reaction containing only Fl-(mt)pre-tRNA^Leu(UUR)^ is marked by closed circles. **B.** Dependence of single turnover rate constant, *k*_obs_, for cleavage of (mt)pre-tRNA^Leu(UUR)^ on the concentration of MRPP3 in the presence of 0.5 µM MRPP1/2. Black square: k*_obs_*measured in the presence of 0.5 µM MRPP1/2, Red square: k*_obs_*measured in the presence of 1 µM MRPP1/2. **C.** Average single turnover cleavage rates at saturating MRPP3 in the presence of 0.4 µM MRPP1/2 for investigated pre-tRNAs. **D.** Average observed single turnover cleavage rates catalyzed by saturating concentrations of PRORP1 and 2 for investigated human pre-tRNAs.

All of the (mt)pre-tRNAs that bind with relatively high affinity to the mtRNase P complex ((mt)pre-tRNA^Leu^,(mt)pre-tRNA^Ile^,(mt)pre-tRNA^Met^) have 
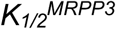
 values, ranging from ~ 390 nM to 930 nM, greater than the measured binding affinities (*K*_D,app_ values in CaCl_2_), suggesting that 
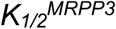
 does not reflect a thermodynamic binding equilibrium under single turnover conditions. The *K*_D,app_ values that we report were measured in CaCl_2_ under conditions that permit binding, but do not promote substrate cleavage. Though unlikely, we cannot rule out the possibility these conditions do not entirely reflect binding of substrates to mtRNase P in MgCl_2_, when cleavage can occur. The *k*_obs_ values for these substrates range between 0.4 - 0.8 min^‒1^ (Figure 3C, Table 1) and correlate well with rate constants previously measured for single-subunit PRORPs cleaving their native substrates (Karasik et al. 2016; Brillante et al. 2016; Howard et al. 2016). In contrast, (mt)pre-tRNAs that have low affinities for mtRNase P ((mt)pre-tRNA^Val^, (mt)pre-tRNASer^UCN^, (mt)pre-tRNA^Ser(AGY)^)) exhibit ~ 100-fold lower single turnover activity (*k*_obs_ ~0.06 min^‒1^) (Figure 3C and Table 1); the cleavage of these substrates was so slow that we were unable to accurately measure 
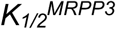
 values. We also considered the possibility that the length of the pre-tRNA 5’ leader or 3’ trailer sequences impacts cleavage by mtRNase P (Supplemental Figure 4). To test this possibility, we measured the rate constant for mtRNase P cleavage of substrates with long length 5’ leaders and 3’ trailers. We found that the length of 3’ and 5’ sequences does not significantly alter the ability of mtRNase P to cleave (mt)pre-tRNA^Ser(UCN)^ and (mt)tRNA^Ile^ constructs. Our findings that the same substrates that bind poorly are also cleaved inefficiently by mtRNase P in single turnover assays (where binding is not the rate-limiting step) suggests that these may not be oriented properly in the bound complex.

As mentioned earlier, we find that most human mitochondrial pre-tRNAs are not bound tightly by *A. thaliana* PRORP1 and PRORP2, therefore we sought to address whether cleavage catalyzed by PRORP1 and 2, as measured by single turnover reaction rates, is also slow for these substrates. In good agreement with our binding data, PRORP1 and PRORP2 processed most human (mt)pre-tRNAs with single turnover rates more than 10-fold slower than that of mtRNase P, except for pre-tRNA^SerUCN^ (Figure 3D, Supplemental Table 2). Pre-tRNA^SerUCN^ is not processed efficiently by the mtRNase P but is cleaved by PRORP1 and 2 with single turnover rate constants comparable to that of their native substrates (~0.3-0.7 min^‒1^) (Figure 3D, Supplemental Table 2). Together with our binding data this suggests that human mitochondrial pre-tRNAs are recognized differently by *A. thaliana* PRORPs and human mtRNase P.

### MRPP3 can enhance the methylation activity of the MRPP1/2 sub-complex

The MRPP1/2 sub-complex was previously shown to methylate *in vitro* transcribed human (mt)pre-tRNAs (Vilardo et al. 2012). Here we measured the single turnover rate constants for MRPP1/2 catalyzing methylation of (mt)pre-tRNAs containing potential m1A9/m1G9 modification sites ((mt)pre-tRNA^Ile^, (mt)pre-tRNA^Leu^ and (mt)pre-tRNA^Val^) (Figure 1, Figure 4A). To determine the rate of methylation, we developed fluorescence primer extension assays for the mtRNase P complex (described in materials and methods, Figure 4B). Additionally, we also assessed if the presence of MRPP3 modulates MRPP1/2 methylation activity. Our data demonstrate that the single turnover rate constants (*k*_obs,meth_) for methylation of (mt)pre-tRNA^Ile^ are 0.13 ± 0.02 min^‒1^ catalyzed by near saturating MRPP1/2 concentrations and that the addition of MRPP3 does not alter the *k*_obs,meth_ value for this substrate (Figure 4C and F,). In comparison, methylation rates for (mt)pre-tRNA^Leu (UUR)^ are a modest 2-fold higher when MRPP3 is present (Figure 4D and F) (0.08 ± 0.01 min^‒1^ in the absence and 0.15 ± 0.03 min^‒1^ in the presence of MRPP3). MRPP1/2 also catalyzed methylation of (mt)-pre-tRNA^Val^, however, the rates are significantly slower (*k_obs,meth_* < 0.005 min^‒1^) compared to the other two pre-tRNA substrates suggesting that there are differences in how MRPP1/2 binds and methylates each pre-tRNA substrate (Figure 4E).

**Figure 4.**
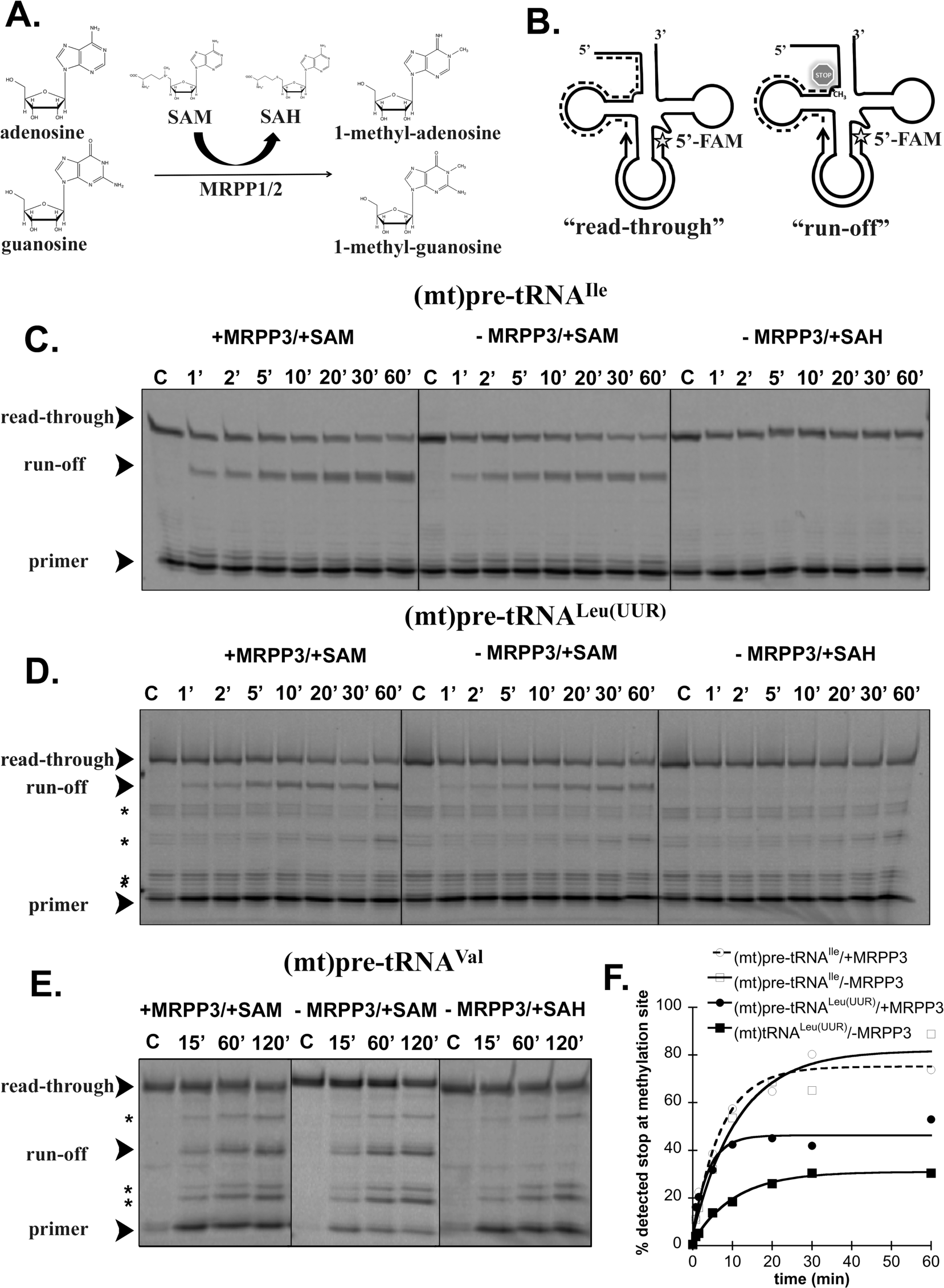
MRPP1/2 methylates human mitochondrial pre-tRNAs.

**A.** Schematic representation of m1A9/m1G9 methylation catalyzed by MRPP1/2 in the presence of S-adenosyl methionine (SAM). SAH=S-adenosyl homocysteine **B.** Scheme for primer extension methylation assays. 5’ fluorescein (5-FAM) labeled primers (black arrow) were designed to base pair with the anticodon loop. In the absence of m1A9/m1G9 methylation, reverse transcriptase (RT) reads all the way to the 5’ end of pre-tRNA (dashed line). When pre-tRNA is methylated at m1A9 or m1G9 position, RT stops at this modification site. **C.**-**E.** Representative gels for methylation of (mt)pre-tRNA^Ile^, (mt)pre-tRNA^Leu^ and (mt)pre-tRNA^Val^ in the presence (first panel) and absence (second panel) of MRPP3. Substitution of SAH for SAM was used as a negative control for methylation (third panel). Black arrows indicate the full-length and methylation stop primer extension products. Asterisks indicate non-specific bands. **F.** Example quantification of the gel-based primer extension methylation assays for (mt)pre-tRNA^Ile^ (open symbols) and (mt) pre-tRNA^Leu^ (closed symbols) in the presence (circles) and absence (squares) of MRPP3.

### SAM is required for mtRNase P to efficiently bind and process some human mitochondrial pre-tRNAs

As discussed above, we found that some human mitochondrial pre-tRNAs are not efficient substrates for the reconstituted mtRNase P complex *in vitro*. However, *in vivo* RNA sequencing experiments suggest that most mitochondrial tRNAs are processed by the mtRNase P complex, including (mt)pre-tRNA^Val^ and (mt)pre-tRNASer^UCN^ (Rackham et al. 2016). This led us to consider whether some substrates, such as (mt)pre-tRNA^Val^ and (mt)pre-tRNASer^UCN^, might require either additional modification steps prior to mtRNase P cleavage, or additional co-factors, such as a protein or small molecule.

As a first step, we investigated if the presence of the methyl donor, SAM, affects the binding affinity of the six (mt)pre-tRNAs used in this study (Figure 1.) We added a high concentration of SAM (25 µM) to our fluorescence anisotropy binding assays and measured (mt)pre-tRNA binding affinities for the MRPP1/2 and mtRNase P complexes (Supplemental Table 3 and 4). We find that the presence of SAM enhances the binding affinity of (mt)pre-tRNA^Val^ for both the MRPP1/2 and the mtRNase P complexes ≥ 15-fold (Figure 5A and B, Supplemental Table 3). Next, we determined if this effect is due to the presence of SAM or methylation of the m1A9 site of (mt)pre-tRNA^Val^. We first measured the (mt)pre-tRNA^Val^ binding affinities for MRPP1/2 and mtRNase P in the presence of the co-factor product of methyl-transfer, SAH, demonstrating an at least ~3 but up to 6-fold decrease in *K*_D,app_ (Figure 5A and B, Supplemental Table 3). To further examine whether methylation is required for the increased binding affinity, we measured the *K*_D,app_ of MRPP1/2 and mtRNase P for a methylation site mutant of (mt)pre-tRNA^Val^ ((mt)pre-tRNA^Val/C9^). The addition of either SAM or SAH enhances the binding affinity of the (mt)pre-tRNA^Val/C9^ mutant to the same extent as for the wild-type substrate (Figure 5A and B, Supplemental Table 3). This suggests that the presence of SAM, but not methylation, is important for (mt)pre-tRNA^Val^ substrate recognition.

**Figure 5.**
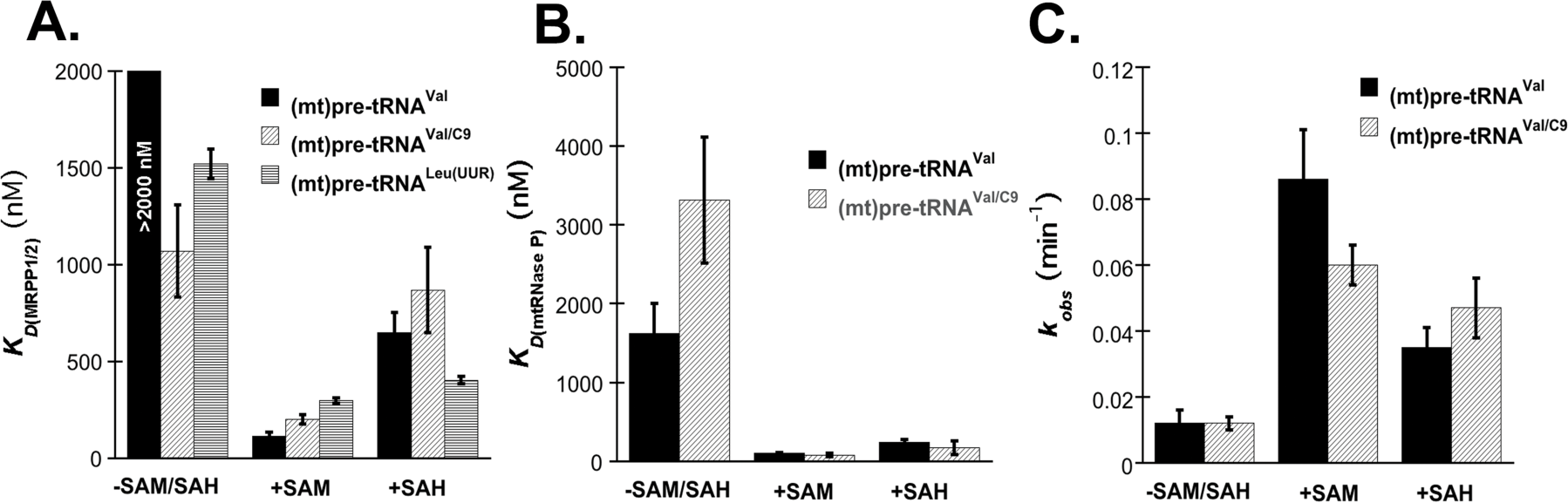
Presence of SAM enhances recognition and 5’ end processing of some mitochondrial pre-tRNAs by mtRNase P.

**A.** Average dissociation constants for (mt)pre-tRNA^Val^ and (mt)pre-tRNA^Leu(UUR)^ for MRPP1/2 in the absence or presence of 25 µM SAM or SAH. **B.** Average dissociation constants for mtRNase P and (mt)pre-tRNA^Val^ in the absence or presence of 25 µM SAM or SAH. **C.** Average observed single turnover cleavage rates for the mtRNase P complex and (mt)pre-tRNA^Val^. Reaction was measured by gel-based cleavage assay in the presence of 625 nM MRPP1/2, 1875 nM MRPP3 and 80 nM labeled pre-tRNA.

In addition, we find that (mt)pre-tRNA^Leu(UUR)^ binds the MRPP1/2 complex ~4-5-fold tighter in the presence of SAM and SAH (Figure 5A), similar to (mt)pre-tRNA^Val^. However, binding affinities for mtRNase P or MRPP1/2 sub-complex for the remaining four substrates are not significantly altered in the presence of SAM (Supplemental Table 4).

We next investigated if the presence of SAM and SAH impact the ability of mtRNase P to cleave the 5’ end of (mt)pre-tRNAs. We find that the single turnover rate constants for the 5’ end cleavage of (mt)pre-tRNA^Val^ by mtRNase P is increased by 3 to 7-fold in the presence of SAH or SAM, respectively (Figure 5C, Supplemental Table 3). Moreover, the single turnover cleavage rates for (mt)pre-tRNA^Val/C9^ catalyzed by mtRNase P exhibit a similar pattern (Figure 5C, Supplemental Table 3). Despite this rate enhancement, it is noteworthy that the rate constant for the processing of (mt)pre-tRNA^Val^ in the presence of SAM is approximately 3 times slower than for any of the other investigated (mt)pre-tRNAs without the presence of SAM (Table 1 and Supplemental Table 3). Addition of SAM or SAH did not enhance the single-turnover rate constants for any of the other (mt)pre-tRNAs that we investigated (Supplemental Table 4). Our results demonstrate that SAM increases the binding efficiency and 5’ end processing catalyzed by mtRNase P for a subset of (mt)pre-tRNAs.

## DISCUSSION

Given the essential nature of human mtRNase P and its direct link to disease it is important to understand how this protein complex functions. Despite its importance, it is mostly unclear how mtRNase P binds and processes its (mt)pre-tRNA substrates. Significantly, how the three components of the mtRNase P interact remains a major question. Here we directly address these critical knowledge gaps by investigating how human mtRNase P binds, cleaves and methylates representative human mitochondrial pre-tRNAs. Our data demonstrate that there is extensive interplay between the mtRNase P components (MRPP1/2/3 and the MRPP1 cofactor SAM) required to recognize, process and methylate human mitochondrial pre-tRNAs. We found that mtRNase P exhibits distinct modes of substrate recognition for individual human mitochondrial pre-tRNAs, in contrast to the homologous single protein component PRORPs that bind and cleave their native substrates more uniformly (Howard et al. 2016).

The mitochondrial genome encodes 22 tRNAs that fall into four different types (0-III) based on their predicted secondary structures. The secondary structures of type 0 (mt)tRNAs are similar to those of canonical nuclear tRNAs, while types I-III vary distinctly. Type I-III (mt)tRNAs differ in the length of their anticodon stems, D‐ and T-loops, or lack the D-loop entirely ((mt)pre-tRNA^Ser(AGY)^ (Rackham et al. 2016))(Watanabe, Suematsu, and Ohtsuki 2014). In our study, we investigated the processing of pre-tRNAs representative of all four categories of mitochondrial pre-tRNAs: (mt)pre-tRNA^Leu^ (type 0), (mt)pre-tRNA^Ser(UCN)^ (type I), (mt)pre-tRNA^Ile^ (type II), (mt)pre-tRNA^Met^ (type II), (mt)pre-tRNA^Val^ (type II) and (mt)pre-tRNA^Ser(AGY)^(type III). These pre-tRNAs are of significant interest because all are hot spots for mutations responsible for a plethora of human mitochondrial diseases (Wortmann et al. 2012; Rossmanith and Karwan 1998; Yan, Zareen, and Levinger 2006; Wang et al. 2011). Additionally, (mt)pre-tRNA^Val^ is of note because it is transcribed between the two mitochondrial rRNAs and is proposed to play an unusual and non-canonical structural role as part of the human mitochondrial ribosome (Amunts et al. 2015; Greber et al. 2015).

Using our selected substrates, we determined both how individual components of the mtRNase P (MRPP3 and MRPP1/2) and the mtRNase P complex bind type 0-III (mt)pre-tRNAs. We find that MRPP3 and MRPP1/2 individually bind most (mt)pre-tRNAs with low affinity (*K_D,app_* values in µM range). However, for (mt)pre-tRNA^Leu^, (mt)pre-tRNA^Ile^, (mt)pre-tRNA^Met^ binding affinities are significantly (*K_D,app_* values in nM range) enhanced when the entire complex is present. This suggests that the different components in the mtRNase complex depend on each other to recognize some substrates. The substrates that bind weakly to mtRNase P include a variety of types (type I (mt)pre-tRNASer^(UCN)^, type II (mt)pre-tRNA^Val^, and type III (mt)pre-tRNASer^(AGY)^). The inability of the complex to bind (mt)pre-tRNA^Ser(AGY)^ was not surprising because this substrate lacks the entire D-loop and stem (Figure 1) and is not predicted to be a substrate of mtRNase P *in vivo* (Rossmanith 1997). However, the weaker binding of (mt)pre-tRNA^Val^ and (mt)pre-tRNASer^(UCN)^ was unexpected since there are no obvious structural or sequence differences that explain the low affinity with mtRNase P.

The efficiency of mtRNase P catalyzing the 5’ end cleavage of particular (mt)pre-tRNAs follows the same pattern as substrate binding; (mt)pre-RNAs that associate tightly are processed 8-20 times faster (single turnover *k*_obs_=0.4 - 0.8 min^‒1^) than substrates that bind weakly (*k*_obs_ ~0.06 min^‒1^). The correlation between weak binding and slow cleavage rates at saturating enzyme concentrations suggests that these pre-tRNA substrates do not bind optimally for cleavage. Notably, all (mt)pre-tRNAs that are recognized and processed efficiently by human mtRNase P are type 0 or II substrates. However, not all type II pre-tRNAs are efficiently processed by mtRNase P, such as pre-tRNA^Val^, suggesting that the ‘type’ of (mt)pre-tRNA structure is insufficient to predict how mtRNase P will recognize any given substrate. Our data indicate that the recombinant mtRNase P does not recognize all substrates equally, leading us to posit that additional factors beyond MRPP1/2 and MRPP3 may be required to ensure that all (mt)pre-tRNAs are efficiently processed in cells.

To gain further insight into the interplay between 5’ end pre-tRNA processing and methylation activities of mtRNase P, we measured the single turnover methylation rates of MRPP1/2 in the presence and absence of MRPP3. These assays were performed using the (mt)pre-tRNAs that possess m1A9/m1G9 methylation sites ((mt)pre-tRNA^Ile^, (mt)pre-tRNA^Leu^, (mt)pre-tRNA^Val^). We find that single turnover rates for methylation were at least 5 times slower than 5’ end processing (Table 1 and Figure 4, e.g. *k_obs,cleav_* ~ 0.8 min^‒1^ and *k_obs,meth_* ~0.13 min^‒1^ for (mt)pre-tRNAIle), suggesting that cleavage may occur prior to methylation. This finding is in agreement with previous studies that assessed (mt)pre-tRNA^Ile^ methylation catalyzed by mtRNase P (Vilardo and Rossmanith 2015). Recent work indicated that 5’ end processing and methylation of (mt)pre-tRNA^Ile^ catalyzed by mtRNase P are carried out by the components of mtRNase P independently, and do not affect one another, further suggesting the structural, but not catalytical role of MRPP1/2 in 5’ end processing (Vilardo et al. 2012; Vilardo and Rossmanith 2015). While our results with (mt)pre-tRNA^Ile^ and (mt)pre-tRNA^Val^ corroborate this suggestion, we find that the presence of MRPP3 modestly increases the rate constant for methylation of (mt)pre-tRNA^Leu^ by MRPP1/2. This implies that the two functions of mtRNase P are not entirely de-coupled, and that there may be cooperation between the 5’ end tRNA processing and methyltransferase functions of mtRNase P for some substrates.

Based on *in vivo* experiments (Rackham et al. 2016; Holzmann et al. 2008), (mt)pre-tRNA^Val^ is predicted to be a substrate of mtRNase P. However, we show that this substrate is not recognized and processed efficiently by recombinant mtRNase P *in vitro*. It has been previously demonstrated that (mt)tRNA^Lys^ requires methylation at the 9^th^ base in order to fully adopt the clover-leaf secondary structure (Helm et al. 1998; Sissler et al. 2004; Motorin and Helm 2010). This result suggested that methylation of (mt)pre-tRNA^Val^ at the 9^th^ position might be necessary for this substrate to fold (and subsequently be processed by mtRNase P), akin to (mt)tRNA^Lys^. Consistent with this model, we observed that the binding and cleavage of (mt)pre-tRNA^Val^ by mtRNase P is significantly enhanced by the addition of SAM (Figure 5). However, similar effects are observed upon either addition of SAH, a SAM analogue that does not function as a cofactor for methylation catalyzed by MRPP1/2, or using the A to C9 methylation site mutant of (mt)pre-tRNA^Val^ (Figure 5). These results suggest that the SAM-mediated effect on (mt)pre-tRNA^Leu^ and (mt)pre-tRNA^Val^ binding affinity and processing by mtRNase P is linked to cofactor binding to MRPP1/2 rather than pre-tRNA methylation. There are several examples in the literature where SAM acts as an effector molecule affecting protein function and activity; SAM is an allosteric regulator of human cystathione-β-synthase and histidine trimethylase EgtD from *M. smegmatis* (McCorvie et al. 2014; Jeong et al. 2014; Lin et al. 2012). Furthermore, in the presence of SAM, a bifunctional restriction enzyme harboring a methyltransferase domain, RM.BpuSI, had increased processing activity without significant methylation activity (Sarrade-Loucheur, Xu, and Chan 2013), similar to mtRNase P acting on (mt)pre-tRNA^Val^. Additionally, a recent study showed an increase in melting temperature for MRPP1 in the presence of SAM indicating a stabilizing effect of the co-factor (Oerum et al. 2018). Based on our data, we propose that the presence of SAM bound to MRPP1 enhances the ability of MRPP1 to bind some pre-tRNA substrates. Our finding also agrees well with the previous observation (Vilardo and Rossmanith 2015) that MRPP1/2 mainly provides structural support for MRPP3.

While we observe that SAM is a key factor for recognition of (mt)pre-tRNA^Val^ by MRPP1/2 and subsequently MRPP3, (mt)pre-tRNA^Ser(UCN)^ is bound to MRPP1/2 efficiently (Table 1) but to mtRNase P poorly. Therefore one possibility is that the recognition of the MRPP1/2-(mt)pre-tRNA^Ser(UCN)^ complex by MRPP3 requires an unidentified mitochondrial factor; however, this needs further testing.

In non-metazoan eukaryotes, the MRPP3-homologous PRORP enzymes are stand-alone proteins able to cleave a wide variety of substrates from different organisms (Gobert et al. 2010; Karasik et al. 2016; Howard, Klemm, and Fierke 2015). To gain insight into potential differences as to how single-protein PRORPs and complex-forming MRPP3 enzymes select and cleave their substrates we assessed the ability of *A. thaliana* PRORP1 and PRORP2 to recognize and process human (mt)pre-tRNAs. Our results suggest that complex-forming protein only RNase Ps recognize their substrates differently than single enzyme PRORPs. Since *A. thaliana* PRORPs recognize the L-shaped tertiary structure of pre-tRNAs (Klemm et al. 2017; Pinker et al. 2017; Imai et al. 2014; Gobert et al. 2013), we propose that mtRNase P may interact with pre-tRNAs by recognizing local structural elements or specific bases within the (mt)pre-tRNAs. Some human mitochondrial tRNAs are known to adopt alternate secondary structures (Lorenz, Lunse, and Morl 2017) and mtRNase P may have conformed to be able to recognize these non-canonical structures.

Taken together, our *in vitro* studies demonstrate that different human mitochondrial pre-tRNAs interact distinctly with mtRNase P and this recognition mode is potentially an important determinant for 5’ tRNA processing. Additionally, we found that some substrates are not efficiently bound and processed by mtRNase P suggesting a role of other factors in mitochondrial 5’end tRNA processing. In line with this, we found that SAM, the cofactor of MRPP1, can enhance substrate recognition and 5’ processing by mtRNAse P. Our data reveal that single subunit PRORPs and human mitochondrial RNase P have distinct modes of pre-tRNA recognition. In addition, our data provide the first evidence that there is an interplay between the 5’ end (mt)pre-tRNA processing and methylation functions of mtRNase P. Together, this work builds on the understanding of the role of MRPP1/2 developed previously (Vilardo et al. 2012) and provides an expanded framework for how the different components of mtRNase P work together to bind, process and methylate mitochondrial pre-tRNAs.

## MATERIALS AND METHODS

### *In vitro* transcription and 5’ end labeling of pre-tRNAs

Pre-tRNAs were prepared as previously described (Karasik et al. 2016; Howard et al. 2012). Briefly, to label the 5’ end of tRNA with fluorescein, we incubated ~200 µL of 5’ GMPS (guanosine-5-monophosphorotioate) labeled *in vitro* transcribed pre-tRNA overnight (in Tris-EDTA buffer of pH 7.2) with 20 µl 45 mM fluorescein (final concentration ~4 mM) at 37 °C and the reaction was purified on a 12% urea-polyacrylamide gel. After gel purification, pre-tRNAs were ethanol precipitated and the resulting pellet was re-suspended in RNase free H_2_O. The concentrations of total and labeled pre-tRNA were measured from absorbance readings at 260 nm using a Nanodrop (Thermo Scientific) spectrophotometer using the following extinction coefficients: 909,100 cm^‒1^ mol^‒1^ for (mt)pre-tRNA^Ile7:0^, 927,300 cm^‒1^ mol^‒1^ for (mt)pre-tRNA^Leu6:0^, 848,600 cm^‒1^ mol^‒1^ for (mt)pre-tRNA^Val7:0^, 838,200 cm^‒1^ mol^‒1^ for (mt)pre-tRNA^Met6:0^, 848,600 cm^‒1^ mol^‒1^ for (mt)pre-tRNA^Ser(UCN)7:0^, 737,900 cm^‒1^ mol^‒1^ for (mt)pre-tRNA^Ser(AGY)7:0^.

### Protein expression and purification

His_6_-Δ95 MRPP3 was cloned into a pMCSG7 vector and transformed into *E. coli* BL21(DE3) cells. Transformed bacteria were grown overnight (5 mL LB media, 100 µg/ml ampicillin, 37 °C (220 rpm)), and used to inoculate 500 mL of TB media with 50 µg/ml ampicillin. Cultures were grown at 37oC to an OD_600_ of ~0.8-1.0, then shifted to 18 °C for 1 hour. 200 µM final concentration of Isopropyl β-D-1 thiogalactopyranoside (IPTG) was added to induce protein expression and the culture was grown overnight (~16 hours) at 18 °C, 220 rpm. Bacterial cells were collected by centrifugation (10,000 x g, 15 minutes), re-suspended in lysis buffer (20 mM MOPS pH 7.8, 1 M NaCl, 1 mM TCEP, 0.2 % (v/v) Tween, 0.1 mg/ml lysozyme and 0.1 mg/ml phenylmethylsulfonyl fluoride). Cells were lysed by sonication, and the resulting suspension was centrifuged at 34,000 x g at 4°C for 1 hour. The supernatant was loaded onto a HisTrap (GE Healthcare) nickel column and protein was eluted with 500 mM imidazole, 20 mM MOPS pH 7.8, 100 mM NaCl and 1 mM TCEP. The purified Δ95 MRPP3 was dialyzed against 20 mM MOPS pH 7.8, 150 mM NaCl, 1 mM TCEP and 15 % glycerol overnight at 4 °C and incubated with TEV protease (1:20 protein:TEV protease ratio) to remove the His tag. His_6_-TEV protease was removed by running the sample through a HisTrap nickel column. Δ95 MRPP3 was then dialyzed against 20 mM MOPS pH 7.8, 1 mM TCEP, 150 mM NaCl and 2 mM EDTA overnight at 4 °C to remove any possible metal contamination, followed by buffer exchange with 20 mM MOPS pH 7.8, 1 mM TCEP and 150 mM NaCl using a desalting column (BioRad). Protein concentration was measured using a Nano spectrophotometer (Thermo scientific) with molecular weight and extinction coefficient values of 56.4 kDa and 82500 cm^‒1^ mol^‒1^, respectively.

Full length MRPP2 and His_6_-Δ39 MRPP1 were cloned and co-expressed as described (Liu 2013). The MRPP1 N-terminal deletion corresponds to the predicted mitochondrial targeting sequence. Initial steps for protein expression and purification, including nickel column purification, histidine tag and TEV removal are identical to those detailed above for Δ95MRPP3, with the following exceptions: the antibiotic used was streptomycin (50 µg/ml) and 10% glycerol was added to each purification buffer. The resulting Δ39 MRPP1/MRPP2 sub-complex was further purified by size exclusion chromatography in 20 mM MOPS pH 7.8, 1 mM TCEP, 100 mM NaCl and 10 % glycerol. Final protein samples were concentrated to 2-4 mg/ml. The ratio of the MRPP1 and MRPP2 proteins in the sub-complex was determined to be 1:4 based on analytical ultracentrifugation experiments. Therefore, protein concentration was measured based on molecular weight and extinction coefficient values of 149 kDa and 79520 cm^‒1^ mol^‒1^. *A. thaliana* PRORP1 and PRORP2 were expressed and purified from *E. coli* as previously described (Howard et al. 2012; Karasik et al. 2016).

### 5’ end cleavage single turnover cleavage assays

(mt)pre-tRNA substrates were re-folded prior to each cleavage reaction. Pre-tRNAs samples were first heated at 95 °C for 3 minutes and then cooled down to room temperature (~10-15 minutes). Cleavage reaction buffer (final: 30 mM MOPS pH 7.8, 150 mM NaCl, 1 mM MgCl_2_, 1 mM DTT) was added and (mt)pre-tRNAs were further incubated for 30 min at room temperature. All single turnover reactions were performed in the cleavage reaction buffer at 28 °C. To initiate a reaction, a mixture of 500 nM Δ39 MRPP1/MRPP2 complex and variable concentrations (100 nM - 2 µM) of Δ95 MRPP3 in cleavage reaction buffer were added to 20 nM fluorescently labeled pre-tRNA. Changes in fluorescence polarization were followed at 28 °C using a ClarioStar plate reader (BMG LabTech). In these assays, we found that the changes in anisotropy observed in the presence of the MRPP1/2 sub-complex and MRPP3 in isolation are small (<1-10%) relative to the significant changes in anisotropy due to cleavage catalyzed by the mtRNase P complex and as such do not impact the rate constants that we measure (Figure 3A). The results of our fluorescence polarization kinetic assays were verified using traditional gel-based cleavage assays (Supplemental Figure 1). For gel-based assays, various time points were obtained by quenching the reaction with the addition of 0.05 % Bromo-Phenol-Blue, 0.05 % Xylene Cyanol dye, 50 % m/v urea and 0.1 M EDTA. The resulting samples were fractionated on a polyacrylamide-urea gel (12-20%) and visualized by direct scanning on a Storm 860 imager. Densitometry was performed with ImageQuant. Data from at least three independent experiments (unless otherwise indicated) were analyzed using Kaleidagraph 4.1.3. Eq 1. was fit to the data to calculate the observed rate constant (*k*) and the standard error from the fitting where *A* is the endpoint, *B* is the amplitude, and *t* is the time. In addition, to obtain the *K_1/2_*value and maximal rate constant (*k_max_*), Eq 2. was fit to the observed rate constants (*k*) determined at different MRPP3 concentrations (while MRPP1/2 kept at a constant 500 nM). The standard error presented is the error of fitting Eq.2. Furthermore, to ensure that N-terminal truncation of Δ95 MRPP3 did not affect single turnover activity of mtRNase P we also reconstituted mtRNase P with Δ45 MRPP3 lacking only the mitochondrial targeting sequence. We found nearly identical single turnover observed cleavage rates for both complexes (Supplemental Table 1).

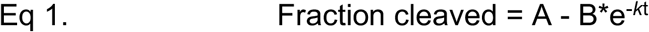

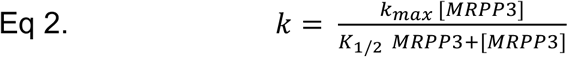

### Fluorescence polarization binding assays

Binding assays were performed in 30 mM MOPS pH 7.8, 150 mM NaCl, 6 mM CaCl_2_, 1 mM DTT at 28 °C and 20 nM of 5’ fluorescein labeled and folded (mt)pre-tRNA. The (mt)pre-tRNA substrates were incubated with increasing concentrations (0-50 µM) of Δ95MRPP3 or MRPP1/2 (0-2 µM) for 5 minutes before changes in anisotropy were measured with ClarioStar plate reader (BMG LabTech). Twenty nM fluorescent pre-tRNA (Fl-pre-tRNA) was titrated by adding increasing concentrations (0-50 µM) of Δ95MRPP3 in the presence of 150 nM Δ39 MRPP1/MRPP2. In some experiments, 25 µM SAM was added to Δ39 MRPP1/MRPP2 and Fl-(mt)pre-tRNA and incubated for 5 minutes before the addition of MRPP3. Baseline anisotropy values measured in the absence of the protein but presence of the Fl-pre-tRNA were extracted from measured values. The dissociation constant (*K_D_*) for PRORPs and apparent dissociation constant (*K*_D,app_) for MRPP1/MRPP2 and mtRNase P and the standard errors were calculated by fitting a binding isotherm (Eq. 3) to the fluorescence polarization data from at least three independent experiments (unless otherwise indicated) using Kaleidagraph 4.1.3 software. In Eq 2, A is the observed anisotropy, A_o_ is the initial anisotropy, ΔA is the total change in anisotropy, P is the concentration of MRPP3, and *K*_D,app_ is the dissociation constant.

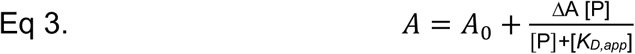

### Primer extension methylation assays

Pre-tRNA was folded as described above before each experiment. Methylation reactions were initiated by mixing the unlabeled pre-tRNA and Δ39MRPP1/MRPP2 to final concentrations of 800 nM of and 5 µM, respectively, in 30 mM MOPS pH 7.8, 150 mM NaCl, 1 mM CaCl_2_ and 1 mM DTT at 28 °C. The observed single turnover methylation rates were measured at varied MRPP1/2 concentrations (2 – 6 µM) for (mt)tRNA^Ile^ and (mt) tRNA^Leu(UUR)^ demonstrating little dependence on the concentration and confirming that 5 µM MRPP1/2 corresponds to a near saturating enzyme concentration (Supplemental Figure 2) To terminate the methylation reaction, 2 µL of enzyme-pre-tRNA reaction mixture was removed, incubated at 95 °C for 30 seconds and then placed on ice. Annealing reactions for primer extensions were set up as follows for each quenched time point. A final volume of a 5 µl annealing reaction contained 1 µl of a quenched methylation reaction and 2 µM of fluorescently labeled primers in 50 mM Tris-HCl (pH 8.1), 30 mM NaCl, and 10 mM DTT. Annealing primer sequences were designed against the anticodon loop for (mt)pre-tRNA^Ile^, (mt)pre-tRNA^Leu^, (mt)pre-tRNA^Val^ and (mt)pre-tRNA^Met^ (6-FAM-CTATTATTTACTCTATC; 6-FAM-CCTCTGACTGTAAAG; 6-FAM-CCTAAGTGTAAGTTGGG; and 6-FAM-CGGGGTATGGGCCCG, respectively). Next, annealing reaction mixtures were incubated at 95°C for 3 minutes followed by cool down to 37°C for 10-15 minutes. As a control, previously folded pre-tRNA was used that was not incubated with the enzymes. 2 µl of annealed primer/(mt)pre-tRNA mixture was added to a final volume of 5 µl reverse transcriptase reaction mixture containing 0.7 U of AMV-Reverse transcriptase (Promega), 1× AMV-RT reaction buffer (Promega) and 0.5 mM dNTP. The reverse transcription reactions were incubated for 1 hour at 37 °C and were subsequently quenched with an equal amount of 0.05 % Bromo-Phenol-Blue, 0.05 % Xylene Cyanol dye, 50 % m/v urea, 0.1 M EDTA and run on 12 % polyacrylamide gel. The gels were scanned in a Storm 860 imager and results were evaluated by densitometry (ImageQuant). As a negative control, we substituted S-adenosyl homocysteine (SAH) for SAM. SAH mimics SAM but lacks the methyl group that is required for the methylation reaction carried out by MRPP1/2. As expected, we did not observe any stops due to methylation in the presence of SAH (Figure 4C-E). Data from at least three independent experiments were analyzed using Kaleidagraph 4.1.3. Eq 1. was fit to the data to calculate the single turnover rate constants (*k_obs, meth_*) and the standard error.

## ACKNOWLEDGEMENTS

We thank Dr. Xin Liu and Dr. Aranganathan Shamuganathan for helping to develop a successful mtRNase P preparation protocol; we also thank the latter for assisting with some of the MRPP1/2 enzyme preparations. We also would like to acknowledge Dr. Nancy Wu and Dr. Xin Liu for supplying Δ45 MRPP3, pre-tRNA^Ser(UCN)36:5^ and pre-tRNA^Ile42:24^. We are grateful to Dr. Bradley Klemm for helpful comments and to Dr. Kristin Koutmou and John Taylor Hosmer-Quint for reading the manuscript. This work was supported by the National Institutes of Health [R01 GM117141 to M.K.] and [R01 GM55387 to C.A.F.]. and by American Heart Association pre-doctoral fellowship [16PRE29890011 to A.K.].

## REFERENCES

Abbott, J. A., C. S. Francklyn, and S. M. Robey-Bond. 2014. ‘Transfer RNA and human disease’, Front Genet, 5: 158.

Amunts, A., A. Brown, J. Toots, S. H. W. Scheres, and V. Ramakrishnan. 2015. ‘Ribosome. The structure of the human mitochondrial ribosome’, Science, 348: 95-98.

Bindoff, L. A., N. Howell, J. Poulton, D. A. McCullough, K. J. Morten, R. N. Lightowlers, D. M. Turnbull, and K. Weber. 1993. ‘Abnormal RNA processing associated with a novel tRNA mutation in mitochondrial DNA. A potential disease mechanism’, J Biol Chem, 268: 19559-64.

Brillante, N., M. Gossringer, D. Lindenhofer, U. Toth, W. Rossmanith, and R. K. Hartmann. 2016. ‘Substrate recognition and cleavage-site selection by a single-subunit protein-only RNase P’, Nucleic Acids Res, 44: 2323-36.

Chatfield, K. C., C. R. Coughlin, 2nd, M. W. Friederich, R. C. Gallagher, J. R. Hesselberth, M. A. Lovell, R. Ofman, M. A. Swanson, J. A. Thomas, R. J. Wanders, E. P. Wartchow, and J. L. Van Hove. 2015. ‘Mitochondrial energy failure in HSD10 disease is due to defective mtDNA transcript processing’, Mitochondrion, 21: 1-10.

Deutschmann, A. J., A. Amberger, C. Zavadil, H. Steinbeisser, J. A. Mayr, R. G. Feichtinger, S. Oerum, W. W. Yue, and J. Zschocke. 2014. ‘Mutation or knock-down of 17beta-hydroxysteroid dehydrogenase type 10 cause loss of MRPP1 and impaired processing of mitochondrial heavy strand transcripts’, Hum Mol Genet, 23: 3618-28.

Gobert, A., B. Gutmann, A. Taschner, M. Gossringer, J. Holzmann, R. K. Hartmann, W. Rossmanith, and P. Giege. 2010. ‘A single Arabidopsis organellar protein has RNase P activity’, Nat Struct Mol Biol, 17: 740-4.

Gobert, A., F. Pinker, O. Fuchsbauer, B. Gutmann, R. Boutin, P. Roblin, C. Sauter, and P. Giege. 2013. ‘Structural insights into protein-only RNase P complexed with tRNA’, Nat Commun, 4: 1353.

Greber, B. J., P. Bieri, M. Leibundgut, A. Leitner, R. Aebersold, D. Boehringer, and N. Ban. 2015. ‘Ribosome. The complete structure of the 55S mammalian mitochondrial ribosome’, Science, 348: 303-8.

Gutmann, B., A. Gobert, and P. Giege. 2012. ‘PRORP proteins support RNase P activity in both organelles and the nucleus in Arabidopsis’, Genes Dev, 26: 1022-7.

Helm, M., H. Brule, F. Degoul, C. Cepanec, J. P. Leroux, R. Giege, and C. Florentz. 1998. ‘The presence of modified nucleotides is required for cloverleaf folding of a human mitochondrial tRNA’, Nucleic Acids Res, 26: 1636-43.

Helm, M., H. Brule, D. Friede, R. Giege, D. Putz, and C. Florentz. 2000. ‘Search for characteristic structural features of mammalian mitochondrial tRNAs’, Rna, 6: 1356-79.

Hodgkinson, A., Y. Idaghdour, E. Gbeha, J. C. Grenier, E. Hip-Ki, V. Bruat, J. P. Goulet, T. de Malliard, and P. Awadalla. 2014. ‘High-resolution genomic analysis of human mitochondrial RNA sequence variation’, Science, 344: 413-5.

Holzmann, J., P. Frank, E. Loffler, K. L. Bennett, C. Gerner, and W. Rossmanith. 2008. ‘RNase P without RNA: identification and functional reconstitution of the human mitochondrial tRNA processing enzyme’, Cell, 135: 462-74.

Howard, M. J., A. Karasik, B. P. Klemm, C. Mei, A. Shanmuganathan, C. A. Fierke, and M. Koutmos. 2016. ‘Differential substrate recognition by isozymes of plant protein-only Ribonuclease P’, Rna, 22: 782-92.

Howard, M. J., B. P. Klemm, and C. A. Fierke. 2015. ‘Mechanistic Studies Reveal Similar Catalytic Strategies for Phosphodiester Bond Hydrolysis by Protein-only and RNA-dependent Ribonuclease P’, J Biol Chem.

Howard, M. J., W. H. Lim, C. A. Fierke, and M. Koutmos. 2012. ‘Mitochondrial ribonuclease P structure provides insight into the evolution of catalytic strategies for precursor-tRNA 5′ processing’, Proc Natl Acad Sci U S A, 109: 16149-54.

Howard, M. J., X. Liu, W. H. Lim, B. P. Klemm, C. A. Fierke, M. Koutmos, and D. R. Engelke. 2013. ‘RNase P enzymes: divergent scaffolds for a conserved biological reaction’, RNA Biol, 10: 909-14.

Imai, T., T. Nakamura, T. Maeda, K. Nakayama, X. Gao, T. Nakashima, Y. Kakuta, and M. Kimura. 2014. ‘Pentatricopeptide repeat motifs in the processing enzyme PRORP1 in Arabidopsis thaliana play a crucial role in recognition of nucleotide bases at TpsiC loop in precursor tRNAs’, Biochem Biophys Res Commun, 450: 1541-6.

Jeong, J. H., H. J. Cha, S. C. Ha, C. Rojviriya, and Y. G. Kim. 2014. ‘Structural insights into the histidine trimethylation activity of EgtD from Mycobacterium smegmatis’, Biochem Biophys Res Commun, 452: 1098-103.

Karasik, A., A. Shanmuganathan, M. J. Howard, C. A. Fierke, and M. Koutmos. 2016. ‘Nuclear Protein-Only Ribonuclease P2 Structure and Biochemical Characterization Provide Insight into the Conserved Properties of tRNA 5′ End Processing Enzymes’, J Mol Biol, 428: 26-40.

Klemm, B. P., A. Karasik, K. J. Kaitany, A. Shanmuganathan, M. J. Henley, A. Z. Thelen, A. J. Dewar, N. D. Jackson, M. Koutmos, and C. A. Fierke. 2017. ‘Molecular recognition of pre-tRNA by Arabidopsis protein-only Ribonuclease P’, Rna.

Klemm, B. P., N. Wu, Y. Chen, X. Liu, K. J. Kaitany, M. J. Howard, and C. A. Fierke. 2016. ‘The Diversity of Ribonuclease P: Protein and RNA Catalysts with Analogous Biological Functions’, Biomolecules, 6.

Li, F., X. Liu, W. Zhou, X. Yang, and Y. Shen. 2015. ‘Auto-inhibitory Mechanism of the Human Mitochondrial RNase P Protein Complex’, Sci Rep, 5: 9878.

Li, R., and M. X. Guan. 2010. ‘Human mitochondrial leucyl-tRNA synthetase corrects mitochondrial dysfunctions due to the tRNALeu(UUR) A3243G mutation, associated with mitochondrial encephalomyopathy, lactic acidosis, and stroke-like symptoms and diabetes’, Mol Cell Biol, 30: 2147-54.

Lin, Y., H. Fan, M. Frederiksen, K. Zhao, L. Jiang, Z. Wang, S. Zhou, W. Guo, J. Gao, S. Li, E. Harrington, P. Meier, C. Scheufler, Y. C. Xu, P. Atadja, C. Lu, E. Li, and X. J. Gu. 2012. ‘Detecting S-adenosyl-L-methionine-induced conformational change of a histone methyltransferase using a homogeneous time-resolved fluorescence-based binding assay’, Anal Biochem, 423: 171-7.

Liu, X., Y. Chen, and C. A. Fierke. 2014. ‘A real-time fluorescence polarization activity assay to screen for inhibitors of bacterial ribonuclease P’, Nucleic Acids Res, 42: e159.

Liu, Xin. 2013. ‘Molecular recognition of inhibitors, metal ions and substartes by Ribonuclease P (Doctoral dissertation)’, Doctoral Dissertation, University of Michigan.

Lorenz, C., C. E. Lunse, and M. Morl. 2017. ‘tRNA Modifications: Impact on Structure and Thermal Adaptation’, Biomolecules, 7.

McCorvie, T. J., J. Kopec, S. J. Hyung, F. Fitzpatrick, X. Feng, D. Termine, C. Strain-Damerell, M. Vollmar, J. Fleming, J. M. Janz, C. Bulawa, and W. W. Yue. 2014. ‘Inter-domain communication of human cystathionine beta‐synthase: structural basis of S-adenosyl-L-methionine activation’, J Biol Chem, 289: 36018-30.

Motorin, Y., and M. Helm. 2010. ‘tRNA stabilization by modified nucleotides’, Biochemistry, 49: 4934-44.

Nickel, A. I., N. B. Waber, M. Gossringer, M. Lechner, U. Linne, U. Toth, W. Rossmanith, and R. K. Hartmann. 2017. ‘Minimal and RNA-free RNase P in Aquifex aeolicus’, Proc Natl Acad Sci U S A, 114: 11121-26.

Oerum, S., M. Roovers, R. P. Rambo, J. Kopec, H. J. Bailey, F. Fitzpatrick, J. A. Newman, W. G. Newman, A. Amberger, J. Zschocke, L. Droogmans, U. Oppermann, and W. W. Yue. 2018. ‘Structural insight into the human mitochondrial tRNA purine N1-methyltransferase and Ribonuclease P complexes’, J Biol Chem.

Ojala, D., J. Montoya, and G. Attardi. 1981. ‘tRNA punctuation model of RNA processing in human mitochondria’, Nature, 290: 470-4.

Pinker, F., C. Schelcher, P. Fernandez-Millan, A. Gobert, C. Birck, A. Thureau, P. Roblin, P. Giege, and C. Sauter. 2017. ‘Biophysical analysis of Arabidopsis protein-only RNase P alone and in complex with tRNA provides a refined model of tRNA binding’, J Biol Chem.

Rackham, O., J. D. Busch, S. Matic, S. J. Siira, I. Kuznetsova, I. Atanassov, J. A. Ermer, A. M. Shearwood, T. R. Richman, J. B. Stewart, A. Mourier, D. Milenkovic, N. G. Larsson, and A. Filipovska. 2016. ‘Hierarchical RNA Processing Is Required for Mitochondrial Ribosome Assembly’, Cell Rep, 16: 1874-90.

Reinhard, L., S. Sridhara, and B. M. Hallberg. 2015. ‘Structure of the nuclease subunit of human mitochondrial RNase P’, Nucleic Acids Res, 43: 5664-72.

Reinhard, L., S. Sridhara, and B. M. Hallberg. 2017. ‘The MRPP1/MRPP2 complex is a tRNA-maturation platform in human mitochondria’, Nucleic Acids Res.

Rossmanith, W. 1997. ‘Processing of human mitochondrial tRNA(Ser(AGY))GCU: a novel pathway in tRNA biosynthesis’, J Mol Biol, 265: 365-71.

Rossmanith, W., and R. M. Karwan. 1998. ‘Impairment of tRNA processing by point mutations in mitochondrial tRNA(Leu)(UUR) associated with mitochondrial diseases’, FEBS Lett, 433: 269-74.

Rossmanith, W., A. Tullo, T. Potuschak, R. Karwan, and E. Sbisa. 1995. ‘Human mitochondrial tRNA processing’, J Biol Chem, 270: 12885-91.

Sanchez, M. I., T. R. Mercer, S. M. Davies, A. M. Shearwood, K. K. Nygard, T. R. Richman, J. S. Mattick, O. Rackham, and A. Filipovska. 2011. ‘RNA processing in human mitochondria’, Cell Cycle, 10: 2904-16.

Sarrade-Loucheur, A., S. Y. Xu, and S. H. Chan. 2013. ‘The role of the methyltransferase domain of bifunctional restriction enzyme RM.BpuSI in cleavage activity’, PLoS One, 8: e80967.

Sen, A., A. Karasik, A. Shanmuganathan, E. Mirkovic, M. Koutmos, and R. T. Cox. 2016. ‘Loss of the mitochondrial protein-only ribonuclease P complex causes aberrant tRNA processing and lethality in Drosophila’, Nucleic Acids Res, 44: 6409-22.

Sissler, M., M. Helm, M. Frugier, R. Giege, and C. Florentz. 2004. ‘Aminoacylation properties of pathology-related human mitochondrial tRNA(Lys) variants’, Rna, 10: 841-53.

Suzuki, T., A. Nagao, and T. Suzuki. 2011. ‘Human mitochondrial tRNAs: biogenesis, function, structural aspects, and diseases’, Annu Rev Genet, 45: 299-329.

Taschner, A., C. Weber, A. Buzet, R. K. Hartmann, A. Hartig, and W. Rossmanith. 2012. ‘Nuclear RNase P of Trypanosoma brucei: a single protein in place of the multicomponent RNA-protein complex’, Cell Rep, 2: 19-25.

Vilardo, E., C. Nachbagauer, A. Buzet, A. Taschner, J. Holzmann, and W. Rossmanith. 2012. ‘A subcomplex of human mitochondrial RNase P is a bifunctional methyltransferase‐‐extensive moonlighting in mitochondrial tRNA biogenesis’, Nucleic Acids Res, 40: 11583-93.

Vilardo, E., and W. Rossmanith. 2015. ‘Molecular insights into HSD10 disease: impact of SDR5C1 mutations on the human mitochondrial RNase P complex’, Nucleic Acids Res, 43: 6649.

Wang, S., R. Li, A. Fettermann, Z. Li, Y. Qian, Y. Liu, X. Wang, A. Zhou, J. Q. Mo, L. Yang, P. Jiang, A. Taschner, W. Rossmanith, and M. X. Guan. 2011. ‘Maternally inherited essential hypertension is associated with the novel 4263A>G mutation in the mitochondrial tRNAIle gene in a large Han Chinese family’, Circ Res, 108: 862-70.

Watanabe, Y., T. Suematsu, and T. Ohtsuki. 2014. ‘Losing the stem-loop structure from metazoan mitochondrial tRNAs and co-evolution of interacting factors’, Front Genet, 5: 109.

Wortmann, S. B., M. P. Champion, L. van den Heuvel, H. Barth, B. Trutnau, K. Craig, M. Lammens, M. F. Schreuder, R. W. Taylor, J. A. Smeitink, R. A. Wevers, R. J. Rodenburg, and E. Morava. 2012. ‘Mitochondrial DNA m.3242G > A mutation, an under diagnosed cause of hypertrophic cardiomyopathy and renal tubular dysfunction?’, Eur J Med Genet, 55: 552-6.

Yan, H., N. Zareen, and L. Levinger. 2006. ‘Naturally occurring mutations in human mitochondrial pre-tRNASer(UCN) can affect the transfer ribonuclease Z cleavage site, processing kinetics, and substrate secondary structure’, J Biol Chem, 281: 3926-35.

